# Nol9 is a Spatial Regulator for the Human ITS2 pre-rRNA Processing Complex

**DOI:** 10.1101/647073

**Authors:** Jacob Gordon, Monica C. Pillon, Robin E. Stanley

## Abstract

The ribosome plays a universal role in translating the cellular proteome. Defects in the ribosome assembly factor Las1L are associated with congenital lethal motor neuron disease and X-linked intellectual disability disorders, yet its role in processing precursor ribosomal RNA (pre-rRNA) is largely unclear. The Las1L endoribonuclease associates with the Nol9 polynucleotide kinase to form the internal transcribed spacer 2 (ITS2) pre-rRNA processing machinery. Together, Las1L-Nol9 catalyzes RNA cleavage and phosphorylation to mark the ITS2 for degradation. While ITS2 processing is critical for the production of functional ribosomes, the regulation of mammalian Las1L-Nol9 remains obscure. Here we characterize the human Las1L-Nol9 complex and identify critical molecular features that regulate its assembly and spatial organization. We establish that Las1L and Nol9 form a higher-order complex and identify the regions responsible for orchestrating this intricate architecture. Structural analysis by high-resolution imaging defines the intricate spatial pattern of Las1L-Nol9 within the nucleolar sub-structure linked with late pre-rRNA processing events. Furthermore, we uncover a Nol9 encoded nucleolar localization sequence that is responsible for nucleolar transport of the assembled LasL-Nol9 complex. Together, these data provide a mechanism for the assembly and nucleolar localization of the human ITS2 pre-rRNA processing complex.

## Introduction

Genes responsible for ensuring the translational capacity of the cell play a prominent role in brain development and function^1–3^. This emerging link highlights the necessity to understand the diverse and dynamic molecular cues orchestrating assembly of the translation machinery^4^. The human ribosome comprises 80 ribosomal proteins and 4 ribosomal RNAs (rRNAs) known as the 18S, 5S, 5.8S and 28S. Ribosomes are assembled through a hierarchical pathway that comprises a series of folding events, integration of ribosomal proteins and precursor rRNA processing^5^. Assembly begins in the nucleolus where RNA polymerase I transcribes the poly-cistronic pre-rRNA which encodes for the 18S, 5.8S and 28S rRNAs separated by two internal transcribed spacers (ITS1 and ITS2) and flanked by two external transcribed spacers (5′-ETS and 3′-ETS)^6^. Efficient and precise removal of these spacers is vital for the functional integrity of the ribosome^6–8^. Correspondingly, there is a growing list of pre-rRNA processing factors associated with motor neuron diseases^6, 9–11^. The pre-rRNA processing factor *LAS1L* is highly expressed in the human cerebellum^12^ and mutations are associated with spinal muscle atrophy with respiratory distress (SMARD) and Wilson-Turner X-linked mental retardation syndrome^13, 14^. Mammalian Las1L is necessary for excision of the ITS2, a pre-rRNA processing event that is paramount for proper maturation of the 60S large ribosomal subunit^15^. Despite its critical role in ribosome assembly, the molecular basis for Las1L regulation in ITS2 processing remains largely unclear.

Las1L associates with the ribosome assembly factor Nol9 to form the ITS2 pre-rRNA processing machinery. Las1L interacts with Nol9 both on and off pre-60S particles^16^. Analogous to Las1L, Nol9 is essential for ITS2 processing and its depletion causes early developmental defects that phenocopy neurodegenerative disease states^16, 17^. Both Las1L and Nol9 are conserved across eukaryotes and together form the ITS2 pre-rRNA processing complex. The mechanism for human ITS2 processing by Las1L-Nol9 is largely unknown, however reconstitution of *Saccharomyces cerevisiae* ITS2 processing has revealed a cascade of catalytic pre-rRNA processing steps^18^. The budding yeast Las1L homolog, Las1, associates with its Nol9 homolog, Grc3^19–21^. The Las1 endoribonuclease initiates ITS2 processing by cleaving a defined site within the ITS2 leaving a 2′-3′-cyclic phosphate and 5′-hydroxyl^8, 22–24^. The resulting 5′-hydroxyl is subsequently targeted by the Grc3 polynucleotide kinase for RNA phosphorylation^8, 25^. The 5′-monophosphate is the molecular signal that coordinates 5′- and 3′-exoribonucleolytic degradation of the ITS2^8, 18, 26, 27^. While these ITS2 processing steps have yet to be confirmed in higher eukaryotes, the strong conservation of the Las1 nuclease motif and Grc3 RNA kinase motifs suggest Las1L is the mammalian endoribonuclease that initially cleaves the ITS2 and Nol9 is the mammalian polynucleotide kinase that phosphorylates the resulting 5′-hydroxyl end of the ITS2 (Fig. 1a)^24, 28^. This is supported by a recent proteomic study confirming the direct interaction between human Las1L-Nol9 and pre-rRNA^29^, as well as earlier work reporting a block in ITS2 excision upon siRNA depletion of mammalian Las1L or Nol9^25, 30, 31^.

**Fig. 1.**
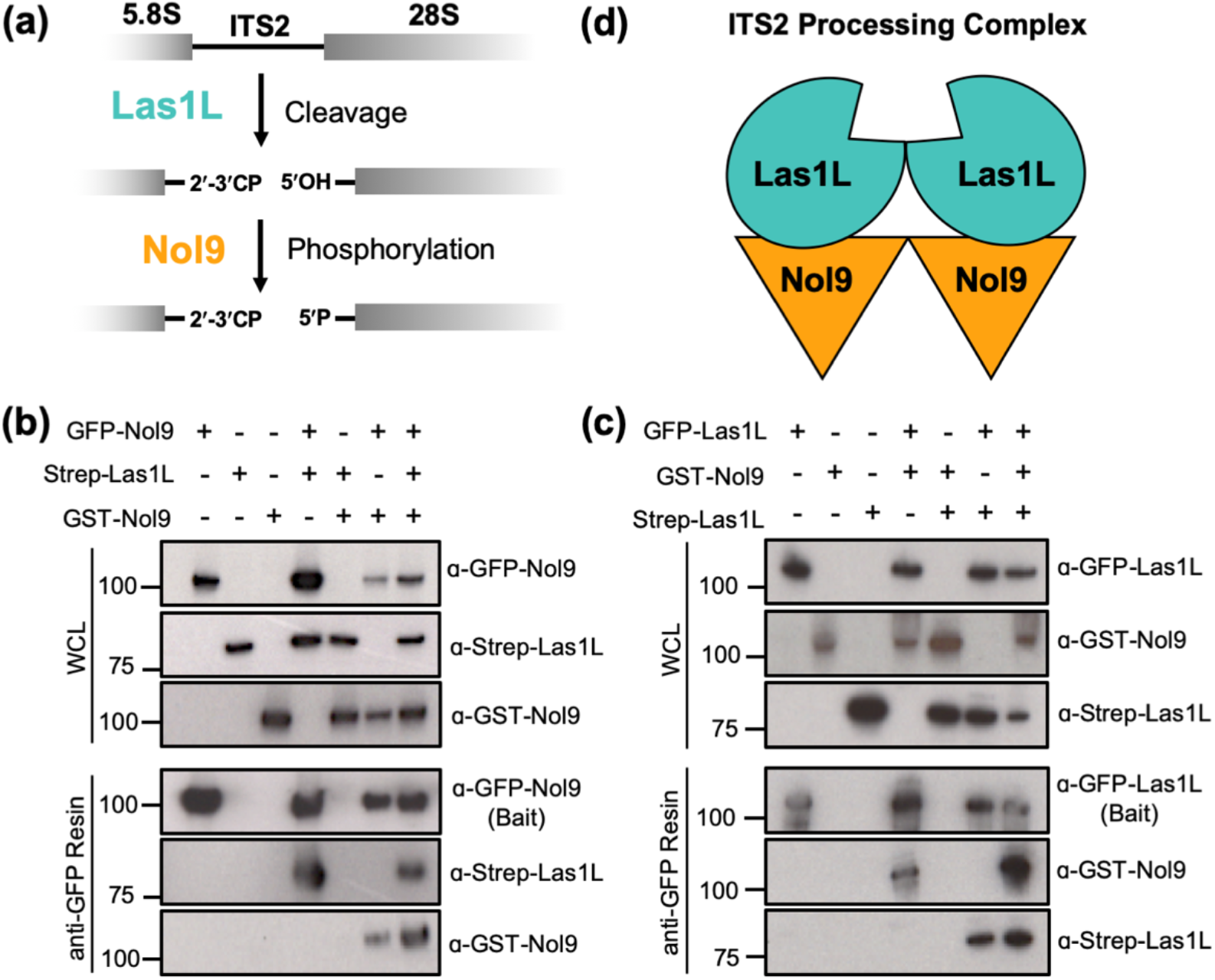
Human Las1L-Nol9 assembles into a higher-order complex. (a) Initial steps of human ITS2 pre-rRNA processing. The Las1L endoribonuclease cleaves the ITS2 at site 4 leaving a 2′-3′ cyclic phosphate (CP) and 5′-hydroxyl (OH). The Nol9 polynucleotide kinase subsequently phosphorylates (P) the 5′-hydroxyl marking the 5′-end of the ITS2 for degradation. (b) Representative co-immunoprecipitation and western blot analysis of transiently expressed GFP-Nol9 (bait) along with GST-Nol9 and Strep-Las1L. Immobilized GFP-Nol9 retains both GST-Nol9 and Strep-Las1L. WCL defines the whole cell lysate. (c) Representative co-immunoprecipitation and western blot analysis of transiently expressed GFP-Las1L (bait) along with Strep-Las1L and GST-Nol9. Immobilized GFP-Las1L retains both Strep-Las1L and GST-Nol9. (d) Model of the higher-order human Las1L-Nol9 ITS2 pre-rRNA processing complex.

Although the overarching principles governing ITS2 processing are likely conserved, there are fundamental differences between lower and higher eukaryotes that undoubtedly necessitate additional layers of regulation. For instance, sequence alignments reveal large insertions within human Las1L and Nol9 that are absent in their yeast counterparts. What influence these additional segments may have on the function and structural architecture of the Las1L-Nol9 complex is unknown. Moreover, the nucleolus is more compartmentalized in higher eukaryotes^30, 32–35^. How human ITS2 pre-rRNA processing is spatially organized across the functionally distinct nucleolar sub-compartments remains undefined. To identify molecular cues important for coordinating mammalian ITS2 processing, we sought to characterize the human ITS2 pre-rRNA processing complex, composed of Las1L-Nol9. In this study, we show Las1L-Nol9 adopts a higher-order assembly that is mediated by molecular features encoded within the C-terminus of both Las1L and Nol9. Using high-resolution confocal microscopy, we map the position of the ITS2 processing complex to reveal its distinct porous-like nucleolar distribution. Structural analysis assigns Las1L-Nol9 to the nucleolar granular component where late pre-rRNA processing events take place. To understand how the ITS2 pre-rRNA processing complex achieves its spatial distribution, we identified a genuine Nol9 nucleolar localization sequence that is responsible for nucleolar transport of Las1L-Nol9. Taken together, Nol9 serves as a critical spatial regulator that imposes sub-cellular organization on the human ITS2 pre-rRNA processing complex.

## Results

### Las1L-Nol9 Assembles into a Higher-Order Complex

To evaluate the structural organization of the human Las1L-Nol9 complex, we transiently transfected HEK 293T cells with plasmids encoding either N-terminally tagged GFP-Nol9, GST-Nol9 or Strep-Las1L. Immunoprecipitation using anti-GFP resin^36^ and western blot analysis confirmed the robust detection of GFP-Nol9 and the absence of nonspecific interactions with the anti-GFP resin (Fig. 1b). Next, we transiently expressed GFP-Nol9 along with Strep-Las1L or GST-Nol9. Co-immunoprecipitation of Strep-Las1L using immobilized GFP-Nol9 recapitulates earlier work that mammalian Las1L associates with Nol9^16^. Interestingly, we also determined that immobilized GFP-Nol9 retains GST-Nol9 (Fig. 1b). This confirms there are multiple copies of the Nol9 polynucleotide kinase in the human Las1L-Nol9 complex. Retention of GST-Nol9 occurs in the absence and presence of transiently expressed Strep-Las1L suggesting Las1L-Nol9 is a mixed population of complexes that harbor either endogenous Las1L or the transiently expressed Strep-tagged Las1L. We confirmed the retention of endogenous Las1L by mass spectrometry from co-immunoprecipitation samples performed with HEK 293T cells transiently expressing GFP-Nol9 and GST-Nol9 (data not shown). This affirms that both endogenous Las1L and transiently expressed Las1L can associate with N-terminally tagged Nol9.

We also performed the reciprocal experiment to explore whether the human Las1L-Nol9 complex harbors multiple copies of the Las1L endoribonuclease. We transiently expressed GFP-Las1L, Strep-Las1L or GST-Nol9 in HEK 293T cells. Immunoprecipitation using anti-GFP resin exclusively retains GFP-Las1L demonstrating the high-selectivity of the GFP affinity support (Fig. 1c). Co-expression of Las1L with Nol9 reveals GFP-Las1L associates with transiently expressed GST-Nol9. This confirms that we can reconstitute human Las1L-Nol9 by immobilizing either the polynucleotide kinase or endoribonuclease component of the complex. Moreover, we show immobilized GFP-Las1L retains Strep-Las1L (Fig. 1c). This reveals the presence of multiple copies of the Las1L endoribonuclease in the human Las1L-Nol9 complex. Since Strep-Las1L is retained in the absence and presence of transiently expressed GST-Nol9, this again suggests there is a mixed population of Las1L-Nol9 where endogenous Nol9 and GST-Nol9 are interchangeable for complex formation. Protein identification by mass spectrometry confirmed the presence of endogenous Nol9 in co-immunoprecipitation samples where GFP-Las1L and Strep-Las1L were transiently expressed (data not shown). Based on this work and previous biophysical characterization of the yeast hetero-tetrameric Las1-Grc3 complex^22^, we propose that human Las1L-Nol9 assembles into a higher-order complex harboring two copies of the Las1L endoribonuclease and two copies of the Nol9 polynucleotide kinase (Fig. 1d). Together, these data demonstrate Las1L-Nol9 higher-order assembly is conserved in humans.

### C-terminal Features of Las1L and Nol9 are Critical for Complex Formation

We next sought to define the molecular features driving assembly of the human Las1L-Nol9 complex. Nol9 harbors a conserved central polynucleotide kinase (PNK) domain that is flanked by N- and C-terminal domains (NTD and CTD) (Fig. 2a). The PNK domain encodes the catalytic motifs necessary for its RNA phosphotransferase activity^25, 28, 37^, whereas the NTD and CTD have no known function. To identify the minimal region of Nol9 that is required to promote its association with Las1L, we engineered and transiently expressed a series of N-terminal GFP-tagged Nol9 truncations and assessed their binding affinity for Strep-tagged Las1L. Truncations to the Nol9 labile N-terminal tail or NTD show comparable Las1L binding as full length Nol9 (Fig. 2b). This suggests the Las1L binding interface is found within the Nol9 PNK and/or CTD. Because it was previously reported that expression of the isolated Nol9 NTD and CTD are unstable^22^, we transiently expressed the Nol9 PNK domain along with Las1L and determined the PNK is insufficient to interact with Las1L. This is in stark contrast to the Nol9 PNK-CTD construct, which shows a clear interaction with Las1L (Fig. 2b) and suggests the Nol9 C-terminus harbors a prominent Las1L binding interface. To confirm that the Nol9 C-terminus is indispensable for Las1L binding, we characterized a Nol9 truncation that encodes the entire Nol9 sequence up to its CTD (residues 1-479). Indeed, the absence of the C-terminus prevents this construct from interacting with Las1L (Fig. 2b). Therefore, we have identified a functional role for the Nol9 C-terminus and assign its activity to driving complex formation with the Las1L endoribonuclease.

**Fig. 2.**
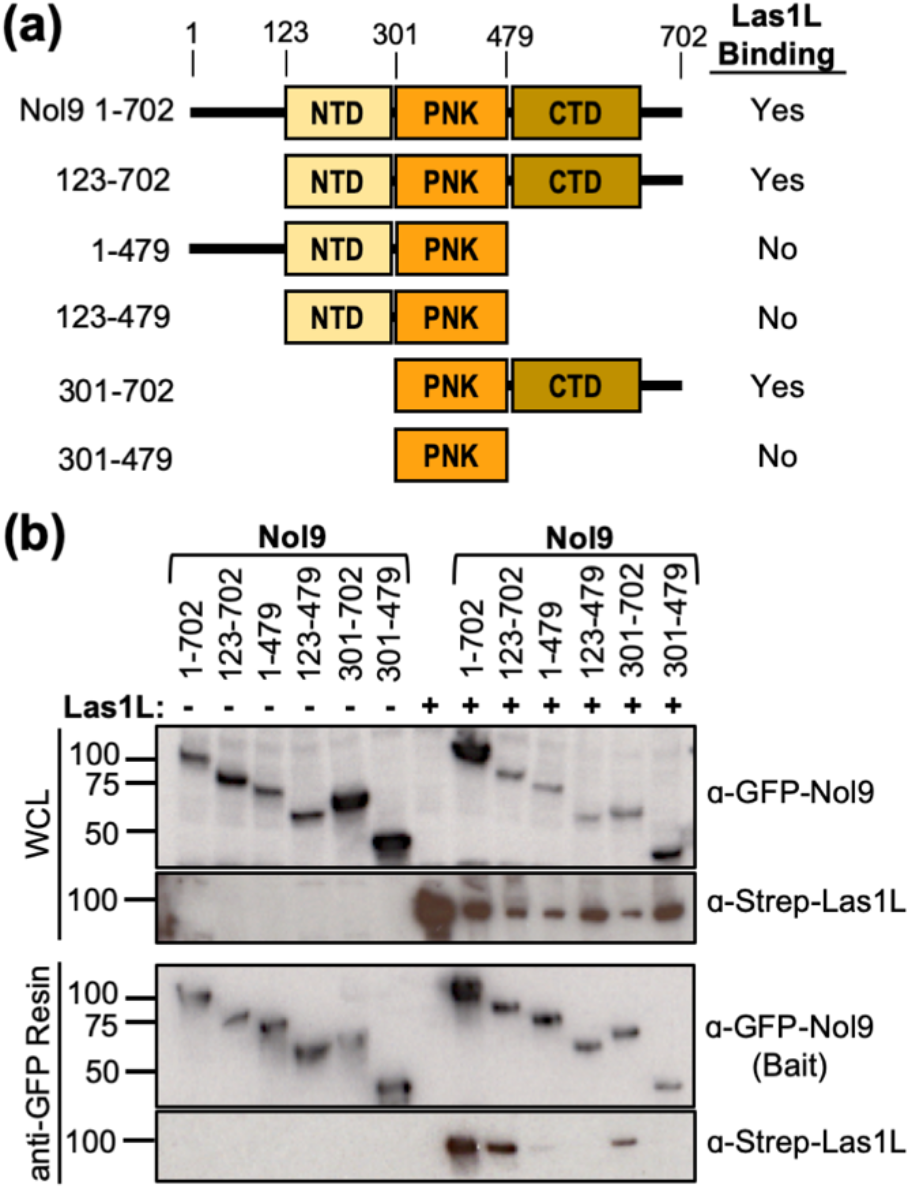
The Nol9 C-terminus is essential for its association with Las1L. (a) Domain architecture of human Nol9. Numbers above the diagram mark the amino acid residue domain boundaries. The yellow box represents the N-terminal domain (NTD), the orange box is the polynucleotide kinase (PNK) domain and the brown box is the C-terminal domain (CTD). (b) Representative co-immunoprecipitation and western blot analysis of transiently expressed GFP-Nol9 truncations (bait) along with full length Strep-Las1L. The Nol9 CTD is necessary to retain Strep-Las1 on the GFP affinity support. WCL defines the whole cell lysate.

We further characterized the Las1L-Nol9 interaction interface using a series of Las1L truncations. Las1L is organized into an N-terminal tail, a Higher Eukaryotes and Prokaryotes Nucleotide-binding (HEPN) domain, a Coiled-coil (CC) domain and a Las1L C-terminal tail (LCT) (Fig. 3a). Human Las1L harbors a characteristic α-helical catalytic HEPN core responsible for its nuclease activity^24^. Unlike other members of the HEPN superfamily, the Las1L/Las1 HEPN domain also contains a functionally undefined N-terminal extension (residues 37-82). To map the features of human Las1L that are required for its association with Nol9, Strep-tagged Las1L truncations and GFP-Nol9 were transiently expressed in HEK 293T cells and co-immunoprecipitated using anti-GFP resin. Since we have previously shown the HEPN core (residues 82-188) is critical for Las1 homodimerization^22^, we characterized Las1L variants that maintained their ability to self-associate. Las1L variants encoding deletions to the N-terminal tail and HEPN extension could still associate with Nol9 demonstrating they are dispensable for complex formation. Moreover, the Las1L HEPN core is insufficient for Nol9 binding since the Las1L variant (residues 1-188) harboring the N-terminal tail along with the HEPN domain did not bind immobilized GFP-Nol9 (Fig. 3b). Conversely, removal of the LCT (residues 614-734) caused a dramatic Nol9-binding defect (Fig. 3b). This demonstrates that the Las1L CC domain is largely dispensable while its LCT harbors critical residues for its association to Nol9. The beginning of the LCT encodes a cluster of conserved hydrophobic residues (Fig. 3a). We asked whether this region of the LCT is important for Las1L-Nol9 complex formation. Interestingly, we could recover Las1L-Nol9 complex formation by extending the Las1L C-terminus to include the LCT hydrophobic patch (residues 614-660) (Fig. 3b). This work defines the LCT hydrophobic patch as a critical region for Nol9 association. This is reminiscent to the yeast ITS2 pre-rRNA processing complex where the hydrophobic C-terminus of Las1 is important for Grc3 association and stability^8, 22^.

**Fig. 3.**
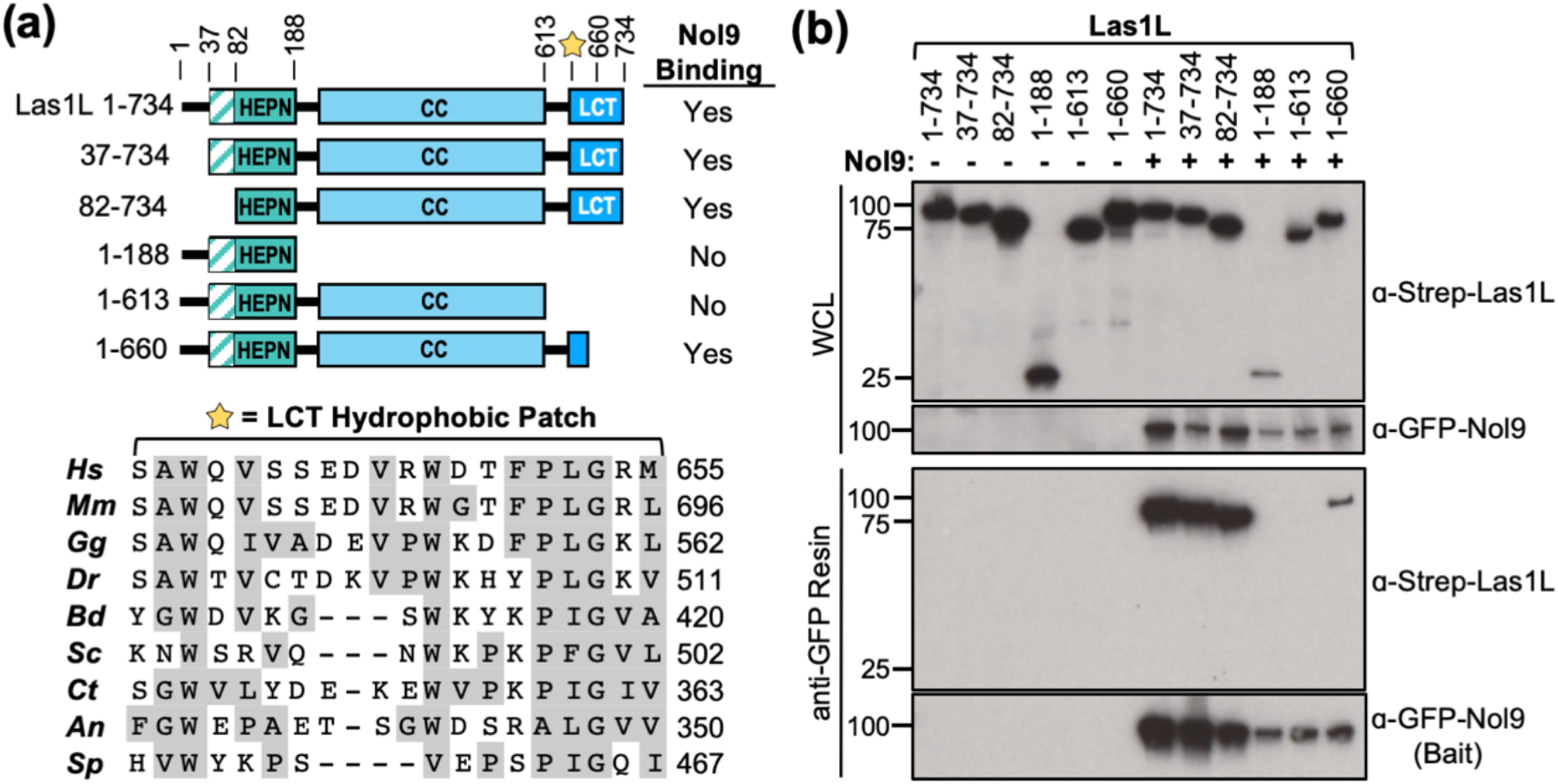
The LCT hydrophobic patch is required for Nol9 binding. (a) Domain architecture of human Las1L. Numbers above the diagram mark the amino acid residue domain boundaries. The striated turquoise box is the HEPN extension, the solid turquoise box is the HEPN core, the light blue box is the coiled-coil (CC) domain and the dark blue box is the Las1L C-terminal tail (LCT). The yellow star marks the location of the conserved LCT hydrophobic patch. Below is the sequence alignment of the conserved LCT hydrophobic patch. Abbreviations are as follows: Hs (*Homo sapiens*), Mm (*Mus musculus*), Gg (*Gallus gallus*), Dr (*Danio rerio*), Bd (*Bactrocera dorsalis*), Sc (*Saccharomyces cerevisiae*), Ct (*Chaetomium thermophilum*), An (*Aspergillus nidulans*), Sp (*Schizosaccharomyces pombe*). (b) Representative co-immunoprecipitation and western blot analysis of transiently expressed Strep-Las1L truncations along with full length GFP-Nol9 (bait). The LCT hydrophobic patch is necessary to retain Strep-Las1L on the immobilized GFP-Nol9 support. WCL defines the whole cell lysate.

### Nol9 Localizes to Regions of Late pre-rRNA Processing

Human Las1L and Nol9 are functionally linked to ITS2 excision^15, 16^, a late pre-rRNA processing event. To precisely map the spatial organization of Las1L-Nol9, we monitored nucleolar localization of N-terminal GFP tagged Nol9 in U2OS cells. Conventional immunofluorescence has demonstrated the nucleolar localization of Nol9^16^, but its detailed spatial organization within the sub-regions of the nucleolus was unknown. To advance our understanding of the sub-nucleolar position of Nol9, we combined confocal laser scanning microscopy with Airyscan to achieve high-resolution imaging^38^. We systematically acquired high-resolution images along different focal planes of the nucleus to render the entire three-dimensional volume of GFP-Nol9 (Fig. 4a). Together, this approach provides high-sensitivity and distinguishes sub-structures within the nucleolus. GFP-Nol9 exclusively localizes to the nucleolus where it displays a distinct porous-like distribution (Fig. 4a). The numerous “pores” are sub-regions void of GFP-Nol9 that are interspersed within extensive areas occupied by GFP-Nol9. This suggests there is a molecular mechanism orchestrating the precise position of Nol9 within the nucleolus.

**Fig. 4.**
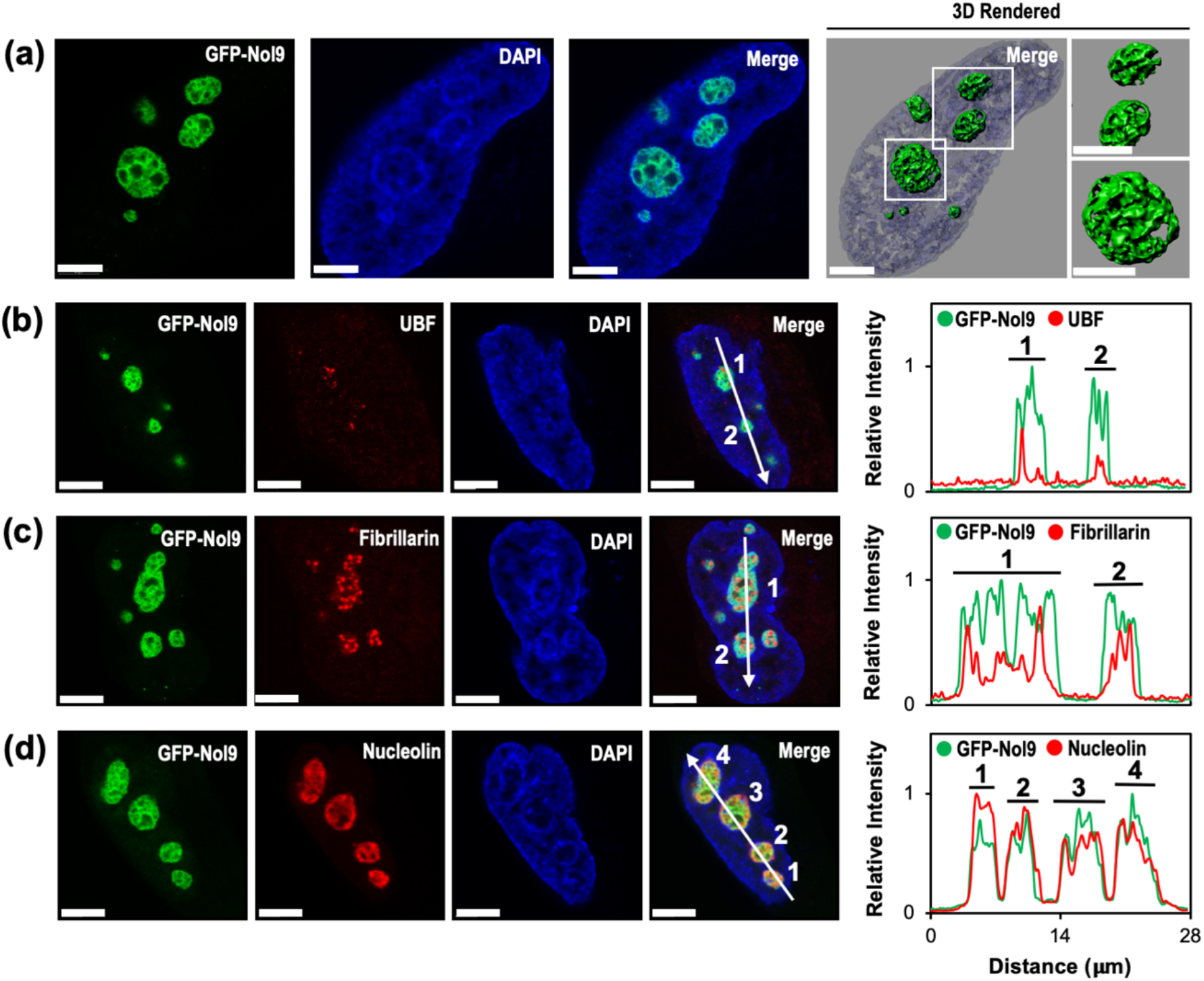
Nol9 localizes to the granular component of the nucleolus. (a) Representative high-resolution image of transiently expressed GFP-Nol9 in fixed U2OS cells. 4′,6-diamidino-2-phenylindole (DAPI, blue) was used to visualize the nucleus. Different focal planes of the nucleus were used to generate a 3D reconstruction (3D Rendered) of GFP-Nol9 within nucleoli. (b-d) Representative high-resolution images of transiently expressed GFP-Nol9 and endogenous nucleolar markers (b) UBF (fibrillar centers), (c) Fibrillarin (dense fibrillar component), and (d) Nucleolin (granular component) in U2OS cells. White arrows mark the path from the images used for the co-localization analysis (graph) whereas the numbers identify each unique nucleolus. Relative intensities of GFP-Nol9 (green) and the nucleolar markers (red) are superimposed and plotted. Scale bars, 4 μm.

We utilized co-localization studies to identify the sub-nucleolar position of Nol9. Human nucleoli are comprised of three distinct sub-regions where each compartment corresponds to a different stage of the pre-rRNA processing pathway^39^. The role of fibrillar centers remains unclear, but has often been associated with rDNA transcription, the dense fibrillar component is linked to early pre-rRNA processing and the granular component is associated with late pre-rRNA processing events^40–42^. Localization of GFP-Nol9 was compared to well-established sub-nucleolar markers, including the Upstream Binding Factor (UBF) for the fibrillar centers, fibrillarin for the dense fibrillar component, and nucleolin for the granular component. Each nucleolar marker displayed a distinct spatial pattern that is characteristic of its respective nucleolar sub-region (Fig.4b-d)^39^. The nucleolar distribution of Nol9 is vastly different from UBF, yet strikingly similar to nucleolin suggesting Nol9 localizes to the granular component (Fig. 4d). Since the spatial organization for fibrillarin is intricate, we quantified the distribution of GFP-Nol9 to each nucleolar marker to generate 2D-plots describing the correlation between their sub-nucleolar localization. The peaks from the intensity profile describing Nol9 localization show a negative relationship with the peaks in the UBF and fibrillarin profiles (Fig. 4b-c). We take this to mean that Nol9 does not have a strong presence in either the fibrillar centers or dense fibrillar component. In contrast, the intensity profiles describing Nol9 and nucleolin localization are largely correlated (Fig. 4d). This work defines the distinct sub-cellular distribution of human Nol9 to regions of the granular component where late pre-rRNA processing events takes place. This is in-line with conventional confocal microscopy showing co-localization of human Las1L with nucleolin^15^.

### Identification of Candidate Nucleolar Localization Sequences in Nol9

Next, we asked how the human Las1L-Nol9 complex achieves its spatial organization within the nucleolus. Las1L nucleolar/nucleoplasm partitioning is sumoylation-dependent^16^, yet the exact molecular mechanism driving Las1L and Nol9 localization to the granular component remains unknown. To determine if either enzyme relies on a nucleolar localization sequence (NoLS), we analyzed the human Las1L and Nol9 primary sequence using the Nucleolar localization sequence Detector (NoD)^43^. This bioinformatic tool aims at identifying candidate NoLSs based on the reoccurring characteristics of validated NoLS signals. Typically, NoLSs are enriched for lysines or arginines, located within a coil, surface accessible and encoded near the N- or C-terminus^44^. Analysis of the Las1L primary sequence did not identify a candidate NoLS while we identified two putative NoLSs in Nol9. The N-terminus of Nol9 contains a putative NoLS (n-pNoLS) 28ILS<u>**RR**</u>P<u>**RRR**</u>LGSL<u>**R**</u>WCG<u>**RRR**</u>L_48_ that is arginine-rich and predicted to lie within a surface exposed coil (Fig. 5a). Furthermore, we identified a putative C-terminal NoLS (c-pNoLS) _681_<u>**R**</u>EPEEAH<u>**K**</u>E<u>**K**</u>PY<u>**RR**</u>P<u>**K**</u>FC<u>**RK**</u>M<u>**K**</u>_702_ that contains both lysine and arginine residues (Fig. 5b). Since both putative NoLSs are promising candidates, we evaluated the functional consequence of deleting each Nol9 pNoLS.

**Fig. 5.**
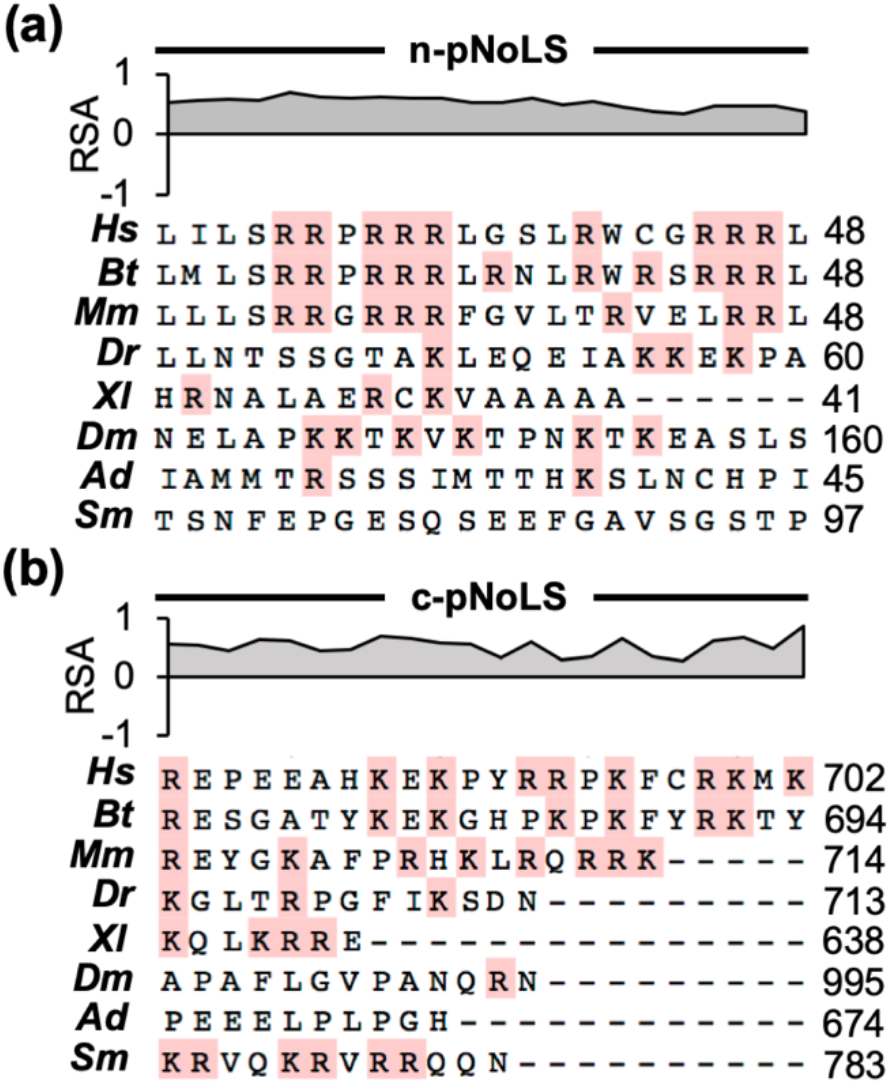
Sequence alignment of candidate Nol9 nucleolar localization signals. Bioinformatic analysis^43^ of the Nol9 primary sequence identified (a) the putative N-terminal nucleolar localization sequence (n-pNoLS) and (b) the putative C-terminal nucleolar localization sequence (c-pNoLS). Red boxes highlight arginine and lysine residues characteristic of NoLS signals. Gray profiles above each sequence alignment displays the predicted relative surface accessibility (RSA) generated by NetSurfP-2.0^58^. Positive RSA values mark residues with a high probability of being surface exposed and negative RSA values identify residues that are likely buried. Abbreviations are as follows: Hs (*Homo sapiens*), Bt (*Bos taurus*), Mm (*Mus musculus*), Dr (*Danio rerio*), Xl (*Xenopus laevis*), Dm (*Drosophila melanogaster*), Ad (*Anopheles darlingi*), Sm (*Stegodyphus mimosarum*).

### Nol9 has an N-terminal Nucleolar Localization Sequence

To determine whether Nol9 relies on an NoLS for its nucleolar localization, we characterized GFP-labeled Nol9 pNoLS variants in U2OS cells using high-resolution imaging. First, we deleted the n-pNoLS (Δn-pNoLS) or c-pNoLS (Δc-pNoLS) from Nol9 and compared its spatial distribution to full length Nol9 (Fig. 6a). Removal of the n-pNoLS largely abrogates Nol9 nucleolar localization and shows homogeneous cellular distribution across the nucleolus, nucleoplasm and cytoplasm (Fig. 6b). There remains a subtle accumulation of GFP-Nol9 Δn-pNoLS in the nucleolus and we attribute this to a sub-population of Las1L-Nol9 complexes that harbor a copy of the endogenous Nol9 protomer. In stark contrast, deleting the Nol9 c-pNoLS shows robust nucleolar localization that is indistinguishable from full length GFP-Nol9 (Fig. 6b). This indicates the c-pNoLS is dispensable for Nol9 nucleolar localization. To validate that the n-pNoLS is sufficient for driving nucleolar localization, we fused the Nol9 n-pNoLS or c-pNoLS sequence to a GFP reporter (Fig. 6a). Transient expression of GFP alone shows homogeneous distribution across the whole cell. Fusing the Nol9 c-pNoLS to the GFP reporter did not significantly alter its broad sub-cellular organization (Fig. 6c). In contrast, GFP fused to the Nol9 n-pNoLS sequence exclusively localizes the GFP reporter to the nucleolus with a porous-like spatial pattern that is reminiscent to GFP-Nol9 localized to the granular component (Fig. 6c). This reveals human Nol9 encodes a genuine NoLS at its N-terminus responsible for directing its position to the granular component of the nucleolus.

**Fig. 6.**
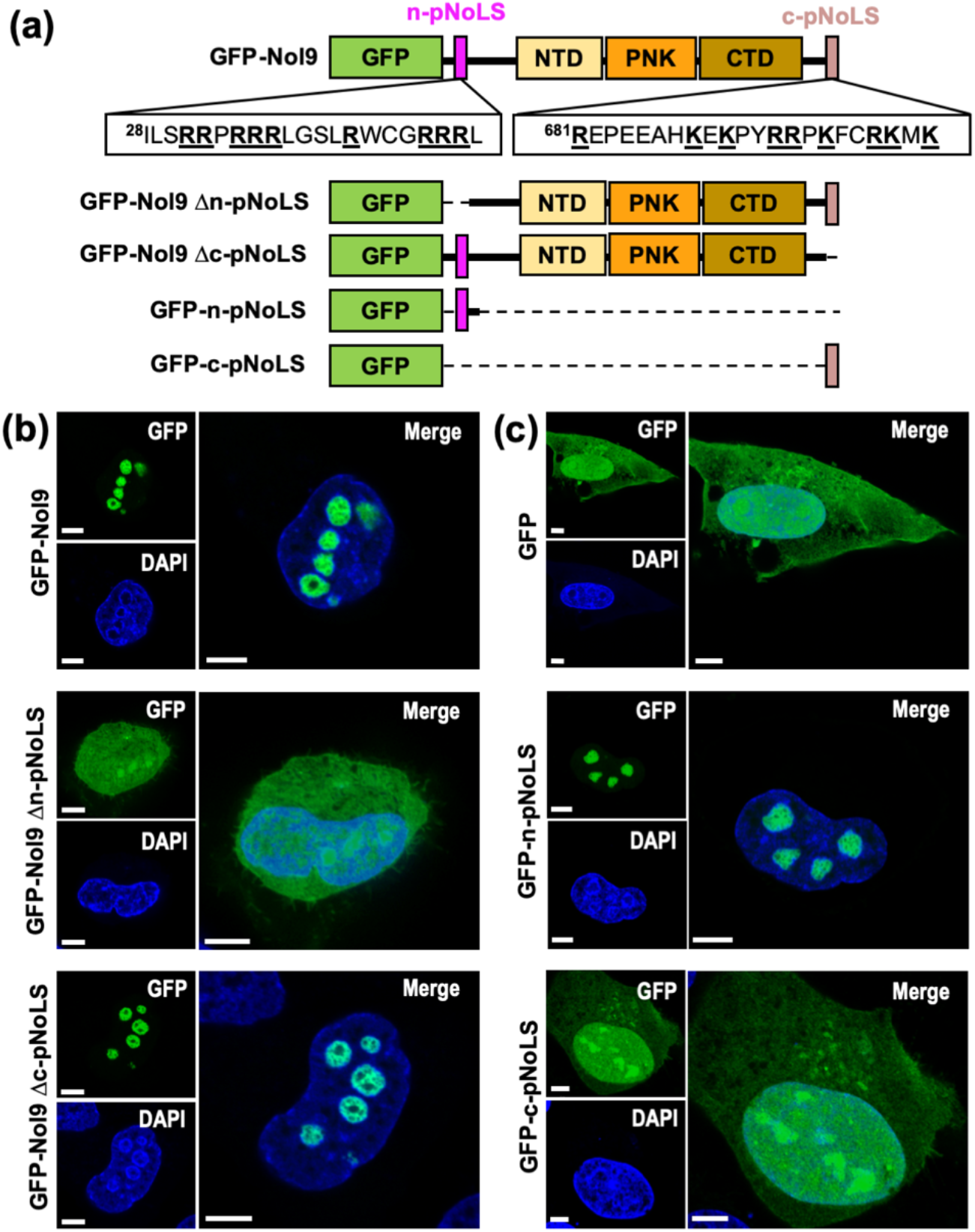
Nol9 n-pNoLS is responsible for Nol9 nucleolar localization. (a) Domain architecture of human Nol9 variants colored as seen in panel 2a. The magenta box represents the putative N-terminal nucleolar localization sequence (n-pNoLS) and the dark beige box represents the putative C-terminal nucleolar localization sequence (c-pNoLS). (b) Representative high-resolution images of transiently expressed GFP-Nol9, GFP-Nol9 Δn-pNoLS and GFP-Nol9 Δc-pNoLS in fixed U2OS cells. (c) Representative high-resolution images of transiently expressed GFP reporter, GFP reporter fused to n-pNoLS and GFP reporter fused to c-pNoLS. 4′,6-diamidino-2-phenylindole (DAPI, blue) was used to visualize the nucleus. Scale bars, 6 μm.

### Nol9 Orchestrates Transport of Las1L into the Nucleolus

Considering Las1L lacks a predicted NoLS, we wondered whether its association with Nol9 may provide a mechanism for nucleolar transport. We performed Nol9 co-localization studies with endogenous Las1L using HeLa cells since the commercial anti-Las1L antibody shows nonspecific binding in U2OS cells. Like its distribution in U2OS cells, GFP-Nol9 has a distinct porous-like pattern in the nucleoli of HeLa cells suggesting it is positioned within the granular component (Fig. 7a). This porous-like pattern is maintained throughout the three-dimensional volume of nucleoli and is largely indistinguishable from the spatial organization of endogenous Las1L (Fig. 7b). Co-localization analysis of several nucleoli produced intensity profiles that confirms a reproducible and robust positive correlation between GFP-Nol9 and endogenous Las1L (Fig. 7c-d). Interestingly, we also observe immunoreactive puncta in the nucleoplasm when using the commercial anti-Las1L antibody in HeLa cells (Fig. 7c). This either reflects non-specific binding of our antibodies or a nuclear sub-population of Las1L. We favor the latter since sumoylation has been shown to regulate Las1L nucleolar/nucleoplasm partitioning^45^. To determine whether the validated Nol9 n-NoLS is required for Las1L nucleolar localization, we monitored endogenous Las1L cellular distribution in transfected HeLa cells expressing GFP-Nol9 Δn-NoLS or GFP-Nol9 Δc-pNoLS. First, we confirmed transient expression of GFP alone does not alter the nucleolar localization of endogenous Las1L (Fig. 8a-b). Similarly, deletion of the Nol9 c-pNoLS had no effect on the spatial distribution of Las1L (Fig. 8c). Conversely, deletion of the validated n-NoLS disrupts the nucleolar localization of both GFP-Nol9 Δn-NoLS and endogenous Las1L in HeLa cells (Fig. 8d). Quantification of their co-localization shows intensity profiles where the homogeneous distribution of Nol9 Δn-NoLS is in sync with endogenous Las1L. Their spatial correlation suggests Las1L is still associated with Nol9 Δn-NoLS, but is no longer capable of positioning itself in the nucleolus. This is in accordance with our co-immunoprecipitation studies where deletion of the Nol9 labile N-terminal tail, which includes the n-NoLS, did not effect Las1L binding (Fig. 2). Thus, Las1L has no intrinsic molecular signal for the nucleolus, but instead associates with Nol9 in the cytoplasm to “piggyback” its way into the nucleolus.

**Fig. 7.**
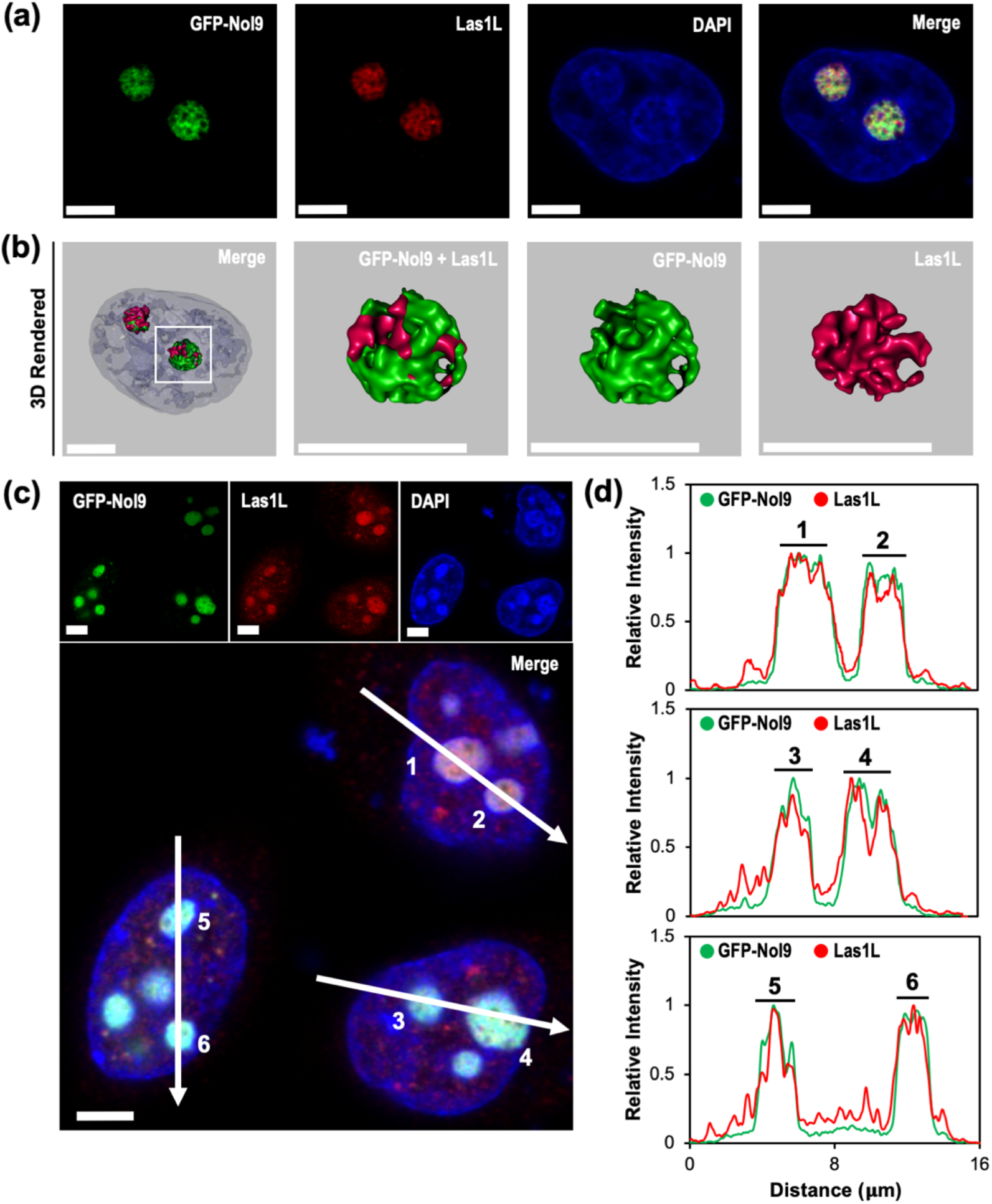
GFP-Nol9 co-localizes with endogenous Las1L. (a) Representative high-resolution image of transiently expressed GFP-Nol9 (green) and endogenous Las1L (red) in fixed HeLa cells. 4′,6-diamidino-2-phenylindole (DAPI, blue) was used to visualize the nucleus. (b) A z-stack was acquired along different focal planes of the nucleus shown in panel a to render a three-dimensional volume of the spatial organization of GFP-Nol9 and endogenous Las1L. (c) Representative high-resolution image of HeLa cells transiently expressing GFP-Nol9. White arrows mark the path for co-localization analysis of GFP-Nol9 and endogenous Las1L. Numbers identify unique nucleoli used for co-localization analysis. (d) Co-localization analysis defines the relative intensities of GFP-Nol9 and endogenous Las1L. Their relative intensity profiles were superimposed and plotted for each nucleolus identified in panel c. Scale bars, 6 μm.

**Fig. 8.**
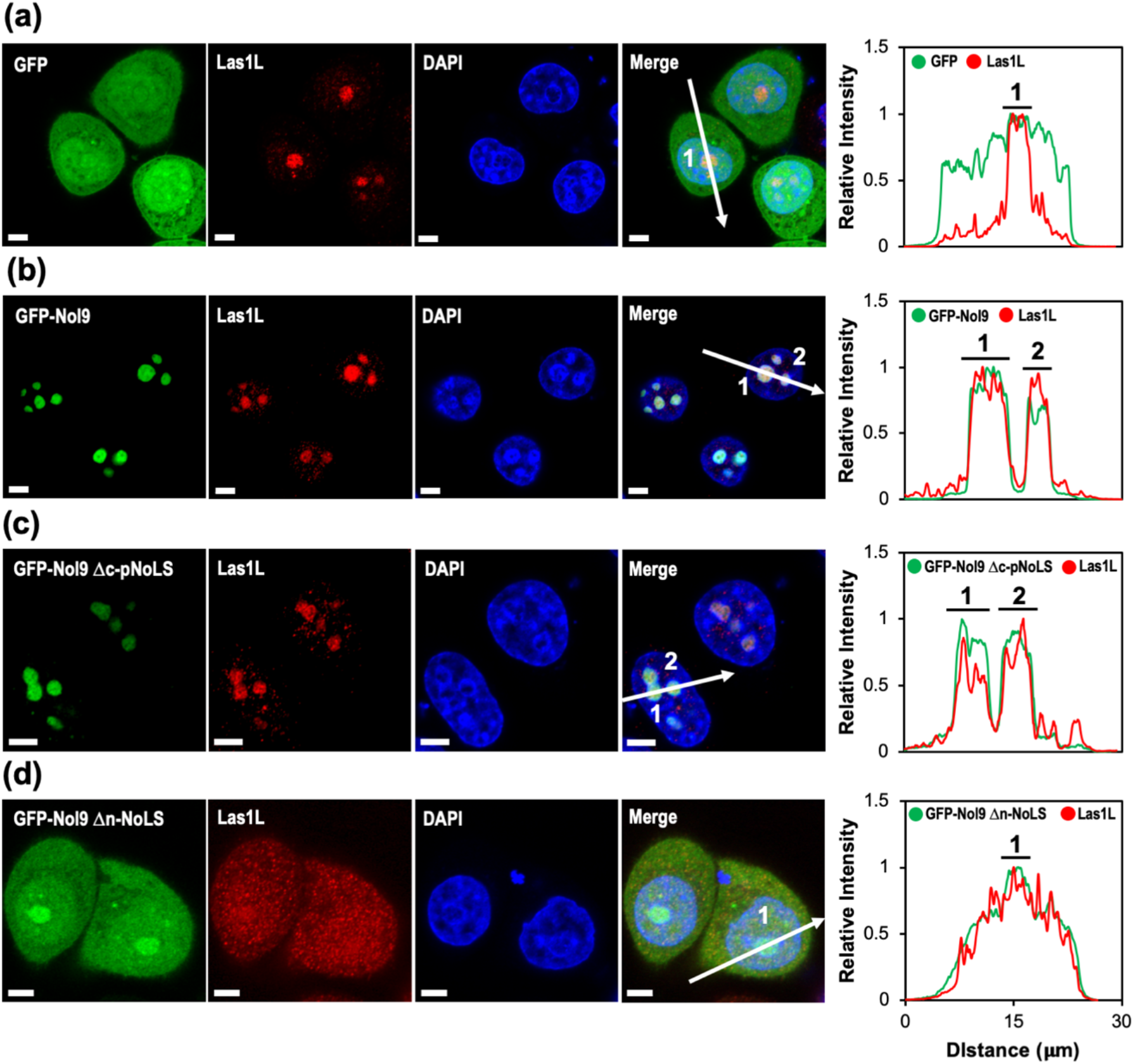
Nol9 transports endogenous Las1L into the nucleolus. Representative high-resolution images of endogenous Las1L and transiently expressed GFP-Nol9 variants in fixed HeLa cells. Endogenous Las1L localization was monitored in cells transiently expressing (a) GFP reporter, (b) GFP-Nol9, (c) GFP-Nol9 Δc-pNoLS and (d) GFP-Nol9 Δn-NoLS. White arrows define the path for co-localization analysis and numbers identify unique nucleoli described in the relative intensity plot shown on the right. Endogenous Las1L relies on the Nol9 n-NoLS for its nucleolar localization. 4′,6-diamidino-2-phenylindole (DAPI, blue) was used to visualize the nucleus. Scale bars, 4 μm.

## Discussion

Here we uncovered the structural organization of the human Las1L-Nol9 pre-rRNA processing complex. We propose the human Las1L-Nol9 complex contains at least two copies of the Las1L endoribonuclease and two copies of the Nol9 polynucleotide kinase (Fig. 9). Members of the HEPN superfamily require dimerization of their HEPN domains for nuclease activation^23, 24, 46–49^. As such, we anticipate the human Las1L HEPN nuclease is a homodimer within the ITS2 pre-rRNA processing complex. This corresponds to previous work revealing the budding yeast Las1 HEPN domain forms a dimer in solution^22^. Following ITS2 cleavage, the associated Nol9 polynucleotide kinase subsequently phosphorylates the resulting 5′-hydroxyl RNA to signal ITS2 degradation^8, 25, 26^. Nol9 belongs to the Clp1/Nol9 PNK family where members share a similar central PNK domain that contains the catalytic motifs necessary for RNA phosphotransferase activity^28, 37, 50, 51^. Interestingly, eukaryotic Clp1 requires a single protomer for RNA phosphorylation^50, 51^. This raises the question as to why there are multiple copies of Nol9 in the ITS2 pre-rRNA processing complex. We speculate that only one catalytically-competent copy of Nol9 is necessary for ITS2 phosphorylation, but two Nol9 protomers are required to scaffold the Las1L homodimer and maintain stability of the complex. This is supported by work performed in yeast showing Las1 relies on its Grc3 binding partner for protein stability and homodimerization^21, 22^. Therefore, this work reveals that Las1L-Nol9 higher-order assembly is not unique to yeast, but a fundamental structural feature of the ITS2 pre-rRNA processing complex that is conserved across the eukaryotic lineage.

**Fig. 9.**
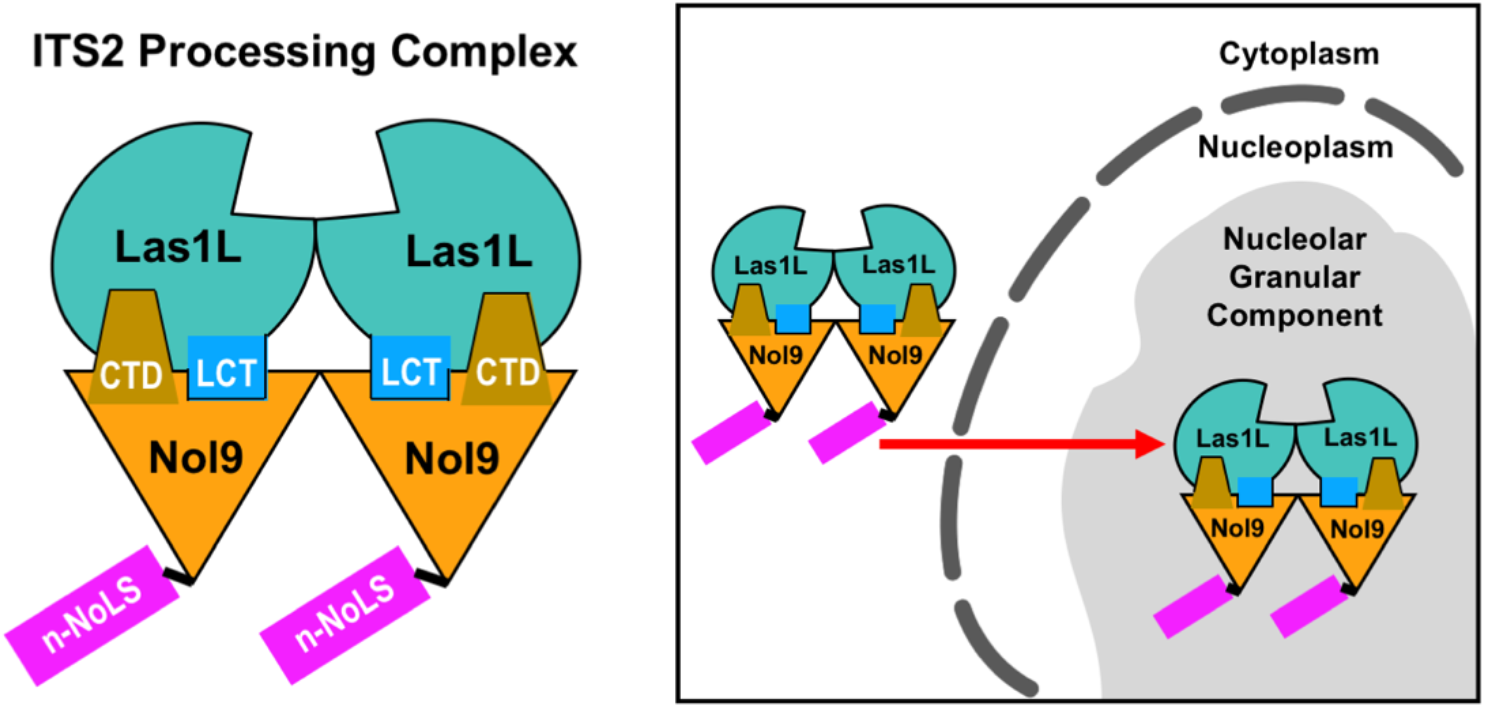
Spatial regulation of the human Las1L-Nol9 complex. The Las1L-Nol9 complex assembles into a higher-order complex. A hydrophobic patch within the Las1L C-terminal tail (LCT) drives the interaction between Las1L and Nol9. The Las1L-Nol9 interface is further supported by the Nol9 C-terminal domain (CTD). Nol9 encodes a labile N-terminal nucleolar localization sequence (n-NoLS) that is responsible for the transport of the assembled Las1L-Nol9 complex into the granular component of the nucleolus where it orchestrates late pre-rRNA processing events, including the removal of the ITS2.

We mapped the molecular features critical for supporting the structural architecture of Las1L-Nol9. Las1L co-immunoprecipitation using Nol9 truncations identified the Nol9 C-terminus as an important region for Las1L binding. The Nol9 C-terminus contains the CTD and a short C-terminal tail that harbors the putative c-pNoLS. Since deleting the Nol9 c-pNoLS did not alter its co-localization with endogenous Las1L (Fig. 8c), this suggests the C-terminal tail is dispensable for Las1L binding. Consequently, we can refine the Las1L binding interface exclusively to the CTD of Nol9. This is an important advancement in our understanding of the Las1L-Nol9 complex because the role of the Nol9/Grc3 CTD was elusive until now. Interestingly, Clp1 also relies on its CTD to mediate a series of protein-protein interactions that are important for its RNA processing activities^52^. This suggests members of the Clp1/Nol9 RNA kinase family may use their CTD as a molecular beacon for the recruitment of RNA processing factors. Moreover, we identified the Las1L hydrophobic patch within the LCT as a minimal region for Nol9 binding. The role of the human LCT as a Nol9-binding factor is consistent with previous characterization of the *S. cerevisiae* Las1-Grc3 complex^8, 22^. While the LCT appears to play a conserved role in Nol9/Grc3 binding, the role of the HEPN domain in complex formation varies across species. We show the human Las1L HEPN domain is insufficient for Nol9 binding, whereas the budding yeast HEPN domain can associate with Grc3^22^. We interpret this to mean that there are variations to the Las1L-Nol9 interaction interface between lower and higher eukaryotes. In fact, Las1L and Nol9 sequence alignments reveal extended termini and insertions that are unique to higher eukaryotes.

This study defines Nol9 as a critical spatial regulator for the human ITS2 pre-rRNA processing complex. We identified putative NoLSs at the N- and C-terminus of Nol9. While the putative c-pNoLS is not a genuine signal, the C-terminal tail is critical for ITS2 cleavage and phosphorylation in yeast suggesting it has a regulatory role^22^. In contrast, the N-terminal NoLS sequence is a bona fide Nol9 nucleolar localization signal. This NoLS transports both Nol9 and Las1L into the nucleolus indicating the complex is first assembled in the cytoplasm (Fig. 9). The presence of a Nol9 NoLS is unlikely to be ubiquitous amongst all homologs since the Nol9 N-terminal extension encoding the NoLS is largely found in higher eukaryotes. Considering this basic N-terminal stretch is absent in primitive eukaryotes, such as invertebrates and fungi, we propose this to be unique to vertebrates (Fig. 5a). Intriguingly, Nol9 sequence alignments suggest the acquisition of the Nol9 NoLS correlates with the transition from a bipartite to tripartite nucleolus^35^. This raises the question as to whether Nol9 acquired the N-terminal NoLS to provide additional regulation for the Las1L-Nol9 complex in species where there is more intricate compartmentalization of the ribosome assembly pathway.

The Nol9 NoLS displays characteristic features that efficiently target the ITS2 pre-rRNA processing complex to the nucleolus. While there is no consensus motif for NoLS signals, recent statistical analysis of validated NoLS targeting motifs have identified general trends. For instance, these signals are often found at polypeptide termini within coils or α-helices that are surface exposed^44^. These characteristic features enhance the accessibility of these motifs to promote their recognition and enhance their effectiveness as a spatial regulator. Since the Nol9 NoLS is found at its N-terminus within a predicted surface exposed coil, we anticipate this targeting motif is constitutively exposed and unobstructed by secondary structure that could dampen its effectiveness. Beyond its accessibility, arginine-rich motifs often equate to stronger NoLS targeting motifs oppose to lysine-rich motifs^53^. This emphasizes that the isoelectric properties of these signals play an important role in determining the potency of these targeting motifs. The Nol9 NoLS does not encode for lysines, but has 9 arginines within a span of 17 residues. Interestingly, previous NoLS characterization has shown that peptides harboring 9 arginine residues display the highest level of nucleolar accumulation^53^. Taken together, the Nol9 NoLS is a strong nucleolar targeting motif. This is in line with its essential role in securing the translational capacity of the cell through pre-rRNA processing of the ITS2. Moreover, the efficient nature of the Nol9 NoLS has the added advantage of safeguarding nonspecific nuclear and cytoplasmic RNAs from aberrant RNA processing by Las1L-Nol9.

Taken together, we establish the conservation of Las1L-Nol9 higher-order assembly in humans and unambiguously identify an NoLS regulatory feature that programs Nol9 to spatially regulate the human ITS2 pre-rRNA processing complex. With the growing list of Las1L and Nol9 binding partners^16, 54^, it will be important to see what influence the Nol9 NoLS may have on the spatial organization of other essential and disease-associated RNA processing factors.

## Materials and Methods

### Expression Vectors of Las1L and Nol9 Variants

*Homo sapiens* Las1L and Nol9 constructs were generated using cDNA provided by GenScript. Full length Las1L (residues 1-734; pMP 062) and Nol9 (residues 1-702; pMP 063) were inserted into a modified eGFP (between EcoRI/KpnI) containing pLEXm vector^55^ using KpnI/XhoI and KpnI/NotI, respectively. Full length Nol9 (pMP 011) was also inserted into pLEXm-GST vector using KpnI/NotI and full length Las1L (pMP 126) was inserted into pCAG-OSF vector^56^ using KpnI/XhoI. Las1L truncations were generated using pMP 126 as a template to amplify Las1L residues 37-734 (pMP 709), residues 82-734 (pMP 708), residues 1-188 (pMP 700), residues 1-613 (pMP 707), residues 1-660 (pMP 701) and inserted into pCAG-OSF vector using KpnI/XhoI. Nol9 truncations were generated using pMP 063 as a template to amplify Nol9 residues 123-702 (pMP 168), residues 1-479 (pMP 167), residues 123-479 (pMP 162), residues 301-702 (pMP 161), residues 301-479 (pMP 163) and inserted into the modified eGFP-pLEXm vector described above using KpnI/NotI. All plasmids were verified by DNA sequencing (GeneWiz).

### Co-immunoprecipitation Assay and Western Blot Analysis

Plasmid transfections were carried out in 40 ml cultures of HEK 293T cells (Invitrogen) that had been adapted to grow in suspension. Cells were transfected with 1 μg/ml of purified plasmid DNA and 2 μg/ml of polyethylenimine (Polysciences Inc.). Cells were harvested 72 hours post transfection and frozen at −80°C until used. Transfected HEK293T cells were lysed in 50 mM Tris pH 8, 100 mM NaCl, 10 mM MgCl_2_, 10% glycerol, 1% Triton X-100, cOmplete EDTA-free protease inhibitor cocktail (Sigma), and 10 units of Benzonase (Sigma) with gentle rocking at 4°C for 45 minutes. Samples were spun at 15,682 × g for 30 minutes at 4°C. Clarified cell lysate was incubated with anti-GFP conjugated resin^36^ for 30 minutes at 4°C. Resin was washed four times with 200 μl of wash buffer (50 mM Tris pH 8, 100 mM NaCl, 10 mM MgCl_2_, 10% glycerol). Samples were resolved by SDS-PAGE and transferred to Immuno-Blot-PVDF membranes (BioRad) using a TE70XP transfer system (Hoefer). Membranes were blocked for 1.5 hours at 22°C using 5% nonfat milk in 1x TBST (20 mM Tris pH 7.6, 150 mM NaCl, 0.1% Tween 20). Antibodies were prepared in 5% nonfat milk, 1% (w/v) BSA and 1x TBST immediately prior to their use. Membranes were incubated in anti-GFP mouse IgG monoclonal primary antibody (Sigma, 11814460001; 1:1000-1:2000), anti-GST mouse IgG monoclonal primary antibody (Santa Cruz, sc-138; 1:1000-1:2000), or anti-Strep mouse IgG monoclonal primary antibody (Sigma, 71590-3; 1:1000-1:2000) overnight at 4°C. Membranes were washed three times in 1x TBST and then incubated for 1 hour at 22°C in anti-mouse IgG HRP conjugate secondary antibody (Sigma, 2834819; 1:1000). Membranes were washed three times in 1x TBST before adding enhanced chemiluminescence detection reagent (GE Healthcare) and developing the blot on film. All co-immunoprecipitation assays were done in triplicate and representative blots are shown in Figures 1, 2, and 3.

### High-resolution Confocal Microscopy

U2OS and HeLa cells (ATCC) were grown on 35-mm glass bottom culture dishes (MatTek Corp.) in DMEM (Invitrogen) supplemented with 10% FBS at 37°C until ~60% confluency. Plasmid DNA (6 μg) was delivered into U2OS cells using FuGENE 6 (Promega) and 2.5 μg of plasmid DNA was delivered into HeLa cells using Lipofectamine 2000 (Invitrogen) according to manufacturer’s instructions. Cells were fixed 1-2 days later in 1x PBS containing 3-4% paraformaldehyde for 20 minutes at 22°C. Cells were then rinsed with 1x PBS and permeabilized with 0.1% Triton X-100 in 1x PBS for 5 minutes followed by two washes with 1x PBS. Cells were incubated for 1 hour at 22°C in blocking buffer containing either 10% normal goat serum in 1x PBS (for endogenous Las1L) or 4% normal goat serum, 1% (w/v) BSA, 0.4% Triton X-100 in 1x PBS (for endogenous Nucleolin, Fibrillarin, UBF). Cells were then washed twice with 1x PBS and then incubated with the appropriate primary antibody for endogenous Las1L (Abcam; ab140656), Nucleolin (Abcam; ab22758), Fibrillarin (Santa Cruz; sc-25397), or UBF (Santa Cruz; sc-9131) overnight at 4°C. Cells were washed three times in 1x PBS and incubated for 1 hour with 10 μg/ml of Alexa Fluor 633 (Invitrogen) in appropriate blocking buffer for Nucleolin, Fibrillarin, and UBF detection or incubated for 45 minutes with 10 μg/ml of Alexa Fluor 568 (Invitrogen) in appropriate blocking buffer for Las1L detection. Cells were washed three times with 1x PBS and incubated for 5 minutes in 300 nM 4′,6-diamidino-2-phenylindole (Sigma) before a final 1x PBS wash. High-resolution images were acquired using a Zeiss LSM 880 inverted confocal microscope with Airyscan^38^ using a 63x/1.4 oil objective. Zen 2012 SP2 (Black Edition) version 11.0 software (Zeiss) was used for data acquisition. Collected images were subjected to the “Airyscan Processing” method using the Zen 2012 SP2 software and Imaris x64 version 9.2.1 (Bitplane) was used for image analysis. Quantification of fluorescent channel profiles were performed using ImageJ/FIJI^57^ and plotted using Excel. For each experiment, over 95% of transfected cells imaged showed the same sub-cellular localization pattern and representative profiles are shown in Fig. 4, 6, 7 and 8.

## Acknowledgements

We would like to thank Dr. Robert Petrovich from the NIEHS Protein Expression Facility for his help with mammalian cell culture. We are also grateful to Dr. Agnes Janoshazi, Charles Tucker, and Erica Scappini from the NIEHS Fluorescence Microscopy and Imaging Center for their technical support. We are indebted to Dr. Jason Williams from the NIEHS Mass Spectrometry Research and Support Group for his assistance with protein identification. We thank Dr. Serena Dudek and Dr. Julie Horton for their critical reading of this manuscript. This work was supported by the US National Institute of Health Intramural Research Program; US National Institute of Environmental Health Sciences (NIEHS; ZIA ES103247 to R.E.S) and the Canadian Institutes of Health Research (CIHR; 146626 to M.C.P).

## Author Contributions

J.G. and M.C.P. designed and constructed vectors for fluorescence microscopy and co-immunoprecipitation experiments. J.G. and M.C.P. performed fluorescence microscopy experiments and J.G. carried out pull-down experiments and western blot analysis. M.C.P. and R.E.S. outlined the study and experiments. J.G., M.C.P. and R.E.S. analyzed the data and prepared the figures. M.C.P. drafted the manuscript which was edited by all the authors.

